# Compost amendments to potato soils enrich yield-associated members of soil microbiomes across the continental US

**DOI:** 10.1101/2025.02.26.640387

**Authors:** Scott A. Klasek, James E. Crants, Kenneth E. Frost, Brenda K. Schroeder, Carl J. Rosen, Linda L. Kinkel

## Abstract

Plant health is regulated by complex consortia of soil microbes with growth-promoting and pathogenic functions. In potato production, various soil management practices are undertaken to boost yields and suppress diseases, but connections between these practices, soil microbiomes, and tuber yields have not been characterized across diverse growing regions. To identify growing practices and microbes associated with increased yields, we established four-year field trials across eight US sites from Oregon to Maine that consisted of controls, fumigations, organic amendments, and mustard incorporations. Soil microbiomes consisted of 16S and ITS amplicon sequences from bacteria and microeukaryotes, respectively. While soil treatments influenced microbiomes differently across all field sites, eukaryotes were more sensitive than bacteria to all treatments. Soil treatments impacted proportions of distinct amplicon sequence variants (ASVs), and associations between ASVs and tuber yields varied within genus-level taxonomy and across field sites. Forty-five ASVs were identified as both treatment-impacted and yield-associated within any field site. Models identified three of thirteen compost amendment scenarios and one of thirteen fumigation scenarios that increased tuber yields by increasing proportions of these taxa within soil microbiomes. These ASVs were not influenced by treatment-associated changes in soil nutrients or organic matter, highlighting complex relationships within specific field sites that require further study to achieve the goal of implementing sustainable, microbiome-informed potato production techniques.

**Importance:** Soil microbes play diverse and interconnected roles in mediating plant health, growth, and disease, but understanding the specifics of how they work and applying them across different agricultural systems remains a challenge. To address this, we amassed a dataset from eight potato field sites across major US growing regions consisting of nearly two thousand soil bacterial and fungal microbiomes paired with soil chemical and tuber yield data. We describe how soil microbiomes are impacted by different soil treatments (compost amendments, chemical fumigation, and mustard incorporation), and identify treatments that boosted yields by up to 23% by increasing proportions of certain bacterial or fungal sequences. Compost amendments affected yield-associated taxa more often than other soil treatments, but these effects varied by rotation length and growing region. Changes to soil chemistry resulting from specific soil treatments did not influence abundances of yield-associated taxa, suggesting that the ways in which they may act to maintain plant vigor are field-specific and complex, calling for further study.

## Introduction

Plants interact with a diversity of soil microbes that play numerous roles in plant fitness, including facilitating nutrient uptake, competing against pathogens, and stimulating host growth through hormonal signaling (1). Changes in plant-associated microbiomes can offer mechanistic insight into these beneficial functions, within the context of plant-microbe-environment interactions (2,3). For these reasons, understanding, applying, and engineering the presence or abundance of specific microbes–and more broadly, the microbiomes they inhabit–has tremendous potential to deliver sustainable agricultural innovations (4,5).

Potatoes are the fifth-most grown crop in the world by yield (6) and play a critical role for global food security based on their high nutrient density and increased cultivation in developing areas (7). Because large-scale cultivation requires intensive soil disturbance and often relies on chemical fumigants like Metam sodium to suppress soil-borne pathogens such as *Verticillium*, alternative strategies that improve soil health while maintaining productivity are of particular interest to growers (8,9). Examples include amending soil with organic matter, which can enhance soil health by delivering nutrients, improving soil physical properties, altering soil microbiomes, suppressing pathogens (10), and even reducing agricultural greenhouse gas emissions (11). In addition, green manures, such as mustards, are sometimes incorporated into soil within potato rotations to reduce soilborne diseases such as Verticillium wilt, black scurf, and common scab; these are thought to act by stimulating pathogen inhibitory activity in soil microbial populations (8,12,13).

Recent studies have identified members of seed tuber microbiomes that are robustly predictive of plant vigor across growing seasons (14), and linked crop rotations and organic matter enrichments with tuber quality and specific microbial taxa (15). Despite this, how soil treatments used in potato cultivation affect relationships between soil bacterial and fungal microbiomes and tuber yields has not been established, particularly across diverse growing regions. We recently described high site-specific variation in the composition of potato soil microbiomes across growing regions of the continental US, which coincided largely with soil physicochemical properties (16).

To investigate whether soil organic amendments, fumigation, or mustard incorporation consistently enriched microbiomes associated with tuber yields across different US regions of potato production, we implemented these treatments across eight US field sites in a four-year field trial. We describe how soil treatments influenced bacterial and eukaryotic potato soil microbiomes across US growing regions, uncover heterogeneous associations between amplicon sequence variants (ASVs) and tuber yields, and use structural equation modeling to identify relationships between organic matter addition, soil chemistry, microbial consortia, and tuber yields. This study provides a benchmark for growers looking to establish sustainable potato production methods, and serves as a starting point for future studies linking cultivation practices (particularly organic matter addition) with growth-associated or disease-suppressive members of soil microbiomes.

## Methods

### Experimental design

Potatoes were grown in two- and three-year rotations from 2019-2022 in eight field sites representing potato growing regions across the continental US, from Oregon to Maine (16). These included six university field sites (OR, CO, MN1, WI, MI, and ME1), one private research site (ID, Miller Research), and one grower field (MN2). Every field site contained six soil and cover cropping treatments corresponding to each rotation length, with four or five replicate plots per treatment. Plots in all fields except CO were arranged in a randomized block design; CO was arranged in a completely randomized design. For the purposes of our analyses, soil treatments were classified as amended with composted or aged manure, conventionally fumigated, or neither (controls). Treatments were also classified based on whether a mustard green manure was tilled back into the soil as a biofumigant at any point in the four-year sequence. Treatments were considered mutually exclusive, except for two irregularities: manure amendments were added to fumigated plots in MI, and manure amendments were confounded with mustard incorporation in MN1. In most cases, local growing regions and practices guided the choice of cultivars and cover crops. Not every field site included a conventional fumigation, organic amendment, and a mustard biofumigant treatment (Fig. 1A). Cover crops varied across field sites, rotations, and treatments; in some cases they were tilled into soils as green manures, but the specific types, sequences, and green manure status of cover crops were not coded into distinct treatment categories. We defined a total of thirty-three management scenarios, each consisting of a soil treatment implemented at a field site under a two- or three-year rotation that corresponded to control treatments growing the same potato cultivar. More detailed treatment information, as well as four-year cropping histories, cultivars grown, rates and types of organic matter and fumigants applied, are provided for all field sites in Supplemental Table S1.

**Fig 1.**
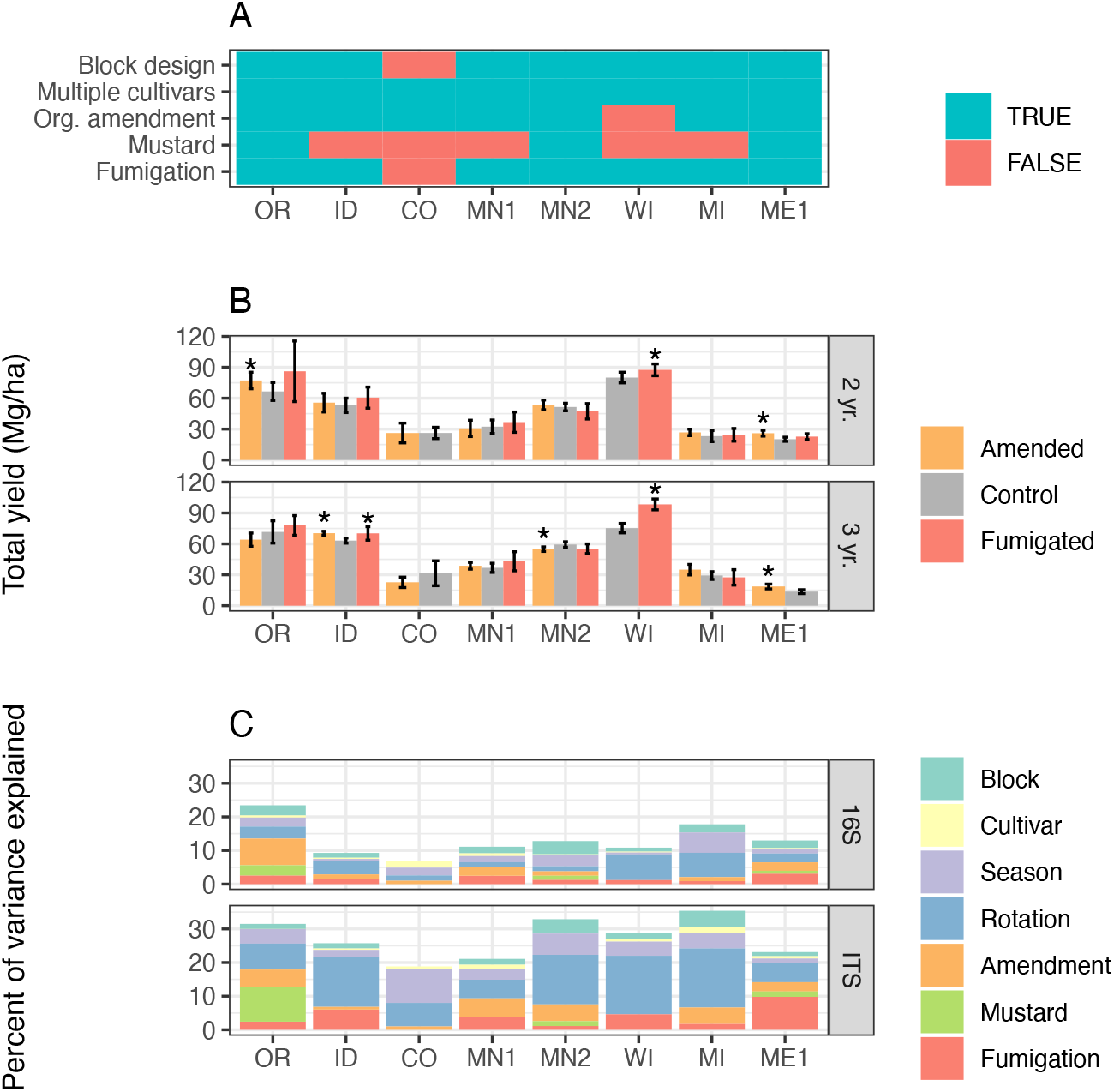
Soil treatments influence tuber yields and microbiome structure. **a** Presence/absence matrix of factors and treatments used in each field site. **b** Total yields, in Mg/hectare, of the most commonly grown cultivar in each field site, separated by soil treatment, site, and rotation length. Error bars indicate 95% confidence intervals, and asterisks show significant differences (p < 0.05) between amended or fumigated treatments relative to controls. **c** Percentages of variance in 16S or ITS microbiome structure attributed to each treatment or factor of the experimental design.

### Soil sampling, DNA extraction, amplicon sequencing, and sequence processing

Potato production soils were sampled and sieved (4.4 mm size) from 456 plots across these eight field sites in 2022 at two seasonal timepoints: at planting and 60 days after planting. DNA was extracted from soils using Qiagen DNeasy PowerSoil Pro kits following manufacturer’s instructions. Further details of sampling, processing, and DNA extraction are described in Klasek et al (16). Amplicons were prepared and sequenced targeting bacterial and microeukaryotic communities with the 16S V3V4 and ITS2 regions, respectively, on an Illumina MiSeq 600 cycle v3 kit (2 × 300 bp reads). Cutadapt v1.18 (17) was used to trim off portions of ITS reads that continued into primer regions of their pairs. Using R (v4.3.2), fastq reads were denoised into amplicon sequence variants (ASVs) with the DADA2 package (18) and taxonomies were assigned using either v138 of the SILVA nonredundant 16S SSU Ref NR99 database (19) or v8.3 of the UNITE database (20,21). The 16S sequences identified as chloroplasts, mitochondria, or eukaryotes were removed, as were the ITS sequences belonging to flowering plants (Anthophyta) or unassigned at the kingdom level. ASV count tables, taxonomy annotations, and sample data were combined into objects using phyloseq v1.46.0 (22). Libraries of either amplicon with less than 10,000 sequences were discarded, leaving 858 16S and 772 ITS soil microbiome samples, with a total of 300,037 16S and 17,214 ITS ASVs. ASVs were assigned unique numbers to ensure consistent identification across different field sites using a custom R function. Microbiome data from 2019 and 2020, as well as sequence prep, sequencing, and amplicon sequence processing steps are described more thoroughly in Klasek et al (16).

### Tuber yields

At the end of the growing season, vines were left to die naturally (ID), killed mechanically (MN1), or killed chemically with one or two applications of diquat (2 pints/acre, all other sites). If repeated, diquat was applied between one and two weeks apart. At least one week after vine death, a total of two rows within each plot (ranging from 5 to 9 m) were harvested by machine from areas that had not been soil sampled earlier in the season. Tubers were weighed and yields were estimated based on row widths (0.86 - 0.91 m). For each site and rotation, two-sided t-tests (alpha = 0.05) were used to determine if yields varied between amended or fumigated treatments relative to controls, with the exception of MI, where amended treatments were compared to fumigated ones because both underwent fumigation. Only the most common cultivar used in each field site was used for these comparisons: Russet Norkotah for OR, ID, and CO; Superior for MI, and Russet Burbank for all other sites. Yields of mustard incorporation treatments were compared directly to corresponding non-mustard treatments.

### Soil chemical and biochemical analyses

A suite of chemical analyses was conducted on soils using previously established techniques (Agvise Laboratories, Benson, MN, USA). Organic matter was measured by loss on ignition at 360°C (23), phosphorus by the Bray and Kurtz spectrophotometric method (24,25), and nitrate-nitrogen by cadmium reduction (26). Ammonium-nitrogen was measured by gas diffusion and conductivity using a KCl solution (Timberline Instruments, Boulder, CO, USA). Soil pH was measured using a 1:1 soil/water slurry (27). Bulk soil respiration was measured by infrared laser spectroscopy upon soil rewetting (28,29). Total bacterial and fungal biomass, derived from phospholipid fatty acid (PLFA) biomolecular signatures, was measured by Ward Laboratories (Kearney, NE, USA).

### Microbiome analyses

#### Variance partitioning analysis of treatments on microbiomes

Proportions of variance in microbiome structure attributable to experimental factors and treatments were determined using version 1.32.2 of the variancePartition R package (30) in R (v4.3.2). For datasets of each amplicon in each field site, ASVs with a mean occupancy below 8.33% were omitted (corresponding to 1 of 12 total treatments), and their counts preserved in a separate column of the count table to maintain compositionality. Count tables were then centered log-ratio (CLR)-transformed. Cultivar, time of sampling (at planting or 60 days after planting), and block number (if applicable) were considered random effects. Fixed effects included rotation length (two levels: two- or three-year) and soil treatment type, wherein logical values corresponded to organic amendment, chemical fumigation, and/or mustard incorporation. The variance attributed to each 16S or ITS ASV from each of these factors was then weighted by ASV mean relative abundance to determine profiles of microbiome-wide variance for each site. ASVs were individually considered to be associated with amended, fumigated, or mustard treatments within particular field sites if the proportion of variance in relative abundance associated with the treatment was ≥ 0.1. This threshold corresponded to the strongest 2.4% and 3.9% of bacterial and eukaryotic ASV-site-treatment associations, respectively.

#### ASV associations with tuber yields

Total tuber yields were imported into sample data of phyloseq objects. Microbiome samples from each field site, rotation length, and sampling time (8 × 2 × 2 = 32 groups total), were analyzed separately. For each group, reads corresponding to ASVs present in < 50% of samples were combined into a single column to preserve compositionality, and counts were then CLR-transformed. Functions within MASS (v.7.3-60.0.1) were used to construct robust linear models of total yields as predicted by CLR-transformed relative abundances of each high-occupancy ASV. Simple linear models were used to obtain summary statistics, including coefficients of determination (R^2^), for each regression. Random subsets of regressions were visually inspected, and ASVs with p-values < 0.05 and multiple R^2^ values ≥ 0.2 were considered to be significantly associated with tuber yields. Forty-five *target ASVs* were identified that both 1) increased in abundance in association with a treatment applied at a field site, and 2) were associated either positively or negatively with tuber yields at the same field site.

#### Modeling soil treatments, ASVs, and yields

Each *management scenario* consisted of a soil treatment at a field site under a two- or three-year rotation that corresponded to a control treatment. In the 16 of 33 scenarios where target ASVs were found, we modeled the treatment and CLR-transformed abundance of a target ASV (or the sum abundance of a combination of such ASVs, where multiple were found) against total yields, using multiple linear regressions as shown in Equation 1.

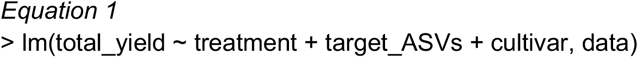

Treatment categories were encoded as dummy variables (0, control; 1, treatment). Different cultivars grown in the same field sites were treated likewise, except in OR and ID data, where only data from the highly predominant Norkotah cultivars were kept. Because ASV associations with tuber yields were specific to the time of soil sampling, models were evaluated for microbiome data collected in 2022 at timepoints where target ASVs were associated with yields (at planting, or 60 days after planting).

Linear models were considered *microbiome-informed* if the increase in abundance of a target ASV (or ASVs) accounted for a significant change in total yield (p < 0.05) that was not directly explained by soil treatment. (Follow-up tests confirmed that target ASV abundances increased in response to soil treatments in all cases).

For scenarios where treatments altered soil chemical constituents (nitrate, ammonium, and phosphate concentrations, pH, % organic matter), we built structural equation models (SEMs) as combinations of linear models using the piecewiseSEM R package v2.3.0 (31) as exemplified in Equation 2.

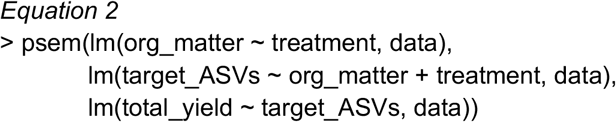

SEMs were considered microbiome-informed if treatments led to increased abundances of a target ASV (or ASVs), which then accounted for a change in total yield that was not directly explained by soil treatment or soil chemical measurements. SEMs were considered valid if p > (higher than) 0.05, corresponding to a failure to reject the model. Cultivars and sampling times were handled as described for linear models. For scenarios where multiple models could be built, all combinations were evaluated, and the one with the lowest AIC (Akaike information criterion) was kept.

## Results

As part of the Potato Soil Health Project (https://potatosoilhealth.cfans.umn.edu/), we characterized 1,824 potato soil microbiomes sampled in 2022 from 456 plots spanning 96 discrete agricultural soil treatments implemented in eight field sites across the continental US from Oregon to Maine. We first evaluated the effects of soil management types (organic amendment, fumigation, or mustard incorporation) on tuber yields and on soil bacterial and eukaryotic microbiomes. Next, we identified amplicon sequence variants (ASVs) associated with tuber yields and specific treatments. Finally, we used analytical models to link soil treatments to ASV-associated changes in tuber yields. Models where treatments changed soil biogeochemistry in ways that influenced yields *independently of the microbiome* are considered beyond the focus of this study.

Total tuber yields in 2022 varied strongly by field site, rotation length, and their interaction (Fig. 1B, two-way ANOVA field site F = 302, p < 2e-16; rotation length F = 9.6, p = 0.002; interaction F = 5.6, p < 4.6e-6). Overall, three-year rotations yielded 3.0 Mg/ha (megagrams, or metric tons per hectare) more than two-year rotations. Across 33 management scenarios that evaluated the direct effects of any soil treatment type, organic amendments significantly increased total tuber yields in 4 of 14 cases (29%), and decreased yields in one case. Fumigation showed a similarly modest impact, increasing yields significantly in 3 of 13 scenarios (23%), with no scenarios decreasing yields. Mustard incorporation, however, did not affect yields in any of six scenarios. Organic amendments in ME1 and fumigations in WI increased yields across both rotation lengths (Fig. 1B), while other interactions between treatments and rotation lengths varied in site-specific ways.

Using linear mixed model-based variance partitioning, we characterized the effects of soil treatments and other experimental factors (cultivar, rotation length, spatial block within the field, and season in which soils were sampled) on the composition of soil bacterial (16S) and eukaryotic (ITS) microbiomes (Fig. 1C). Overall, individual factors explained 0 to 17.4% of bacterial microbiome variance within and across field sites, and collectively explained a mean of 14% of the variance among bacterial microbiomes. For eukaryotic microbiomes, within which 89% of reads were classified to fungi, 28% of variance was explained by these factors. Organic amendment, fumigation, and mustard treatments showed particularly strong effects in some field sites (fumigation in ME1 ITS; mustard in OR ITS; amendment in OR 16S), but not others, and together explained 1.6% more of the variance in the composition of eukaryotic microbiomes as opposed to bacterial ones (p = 0.049, F = 4.22, one-way ANOVA, type of treatment not significant). Rotation length was the factor that explained the most variance in both bacterial (3.9%) and eukaryotic (11.4%) microbiome composition across field sites, while in contrast, potato cultivar explained a mean 0.6% of variance in the structure of each microbiome type.

We applied the same variance partitioning method to individual amplicon sequence variants (ASVs), to discern which taxa were most responsive to soil treatments. As described above, the relative amount of variance from each soil treatment and experimental factor was determined for each ASV. We defined *treatment-associated ASVs* as those for which any soil treatment explained 10% or more of their variance in relative abundance across all plots and both sampling times within a field site. Using this definition, equal proportions of 16S and ITS ASVs (12.5% and 12.6% respectively) were responsive to any treatment at any site. While organic amendment, fumigation, and mustard incorporation affected abundances of different soil genera in varying proportions, all soil treatments imparted a wide range of influence on distinct ASVs belonging to the same bacterial or eukaryotic genus, from less than 1% to over 40% of variance ascribed to treatment (Fig. 2). This suggests that soil treatments have highly variable effects on different members of the same genus. Even for a soil treatment applied across plots within the same field, we noticed a wide range in impact: in ME1, fumigation accounted for less than 1% to 56% of the variance in the relative abundances of ASVs from the fungal genus *Solicoccozyma*, while in OR, organic amendment accounted for anywhere from 5 to 72% of the variance in *Novibacillus* ASV relative abundances (Fig. 2). These single-site findings suggest that the highly variable impacts of soil treatment on different members of a single microbial genus is not simply a consequence of soil chemistry or climate, and may instead reflect differential responses to selection among species, strains, or ASVs within the same genus.

**Fig 2.**
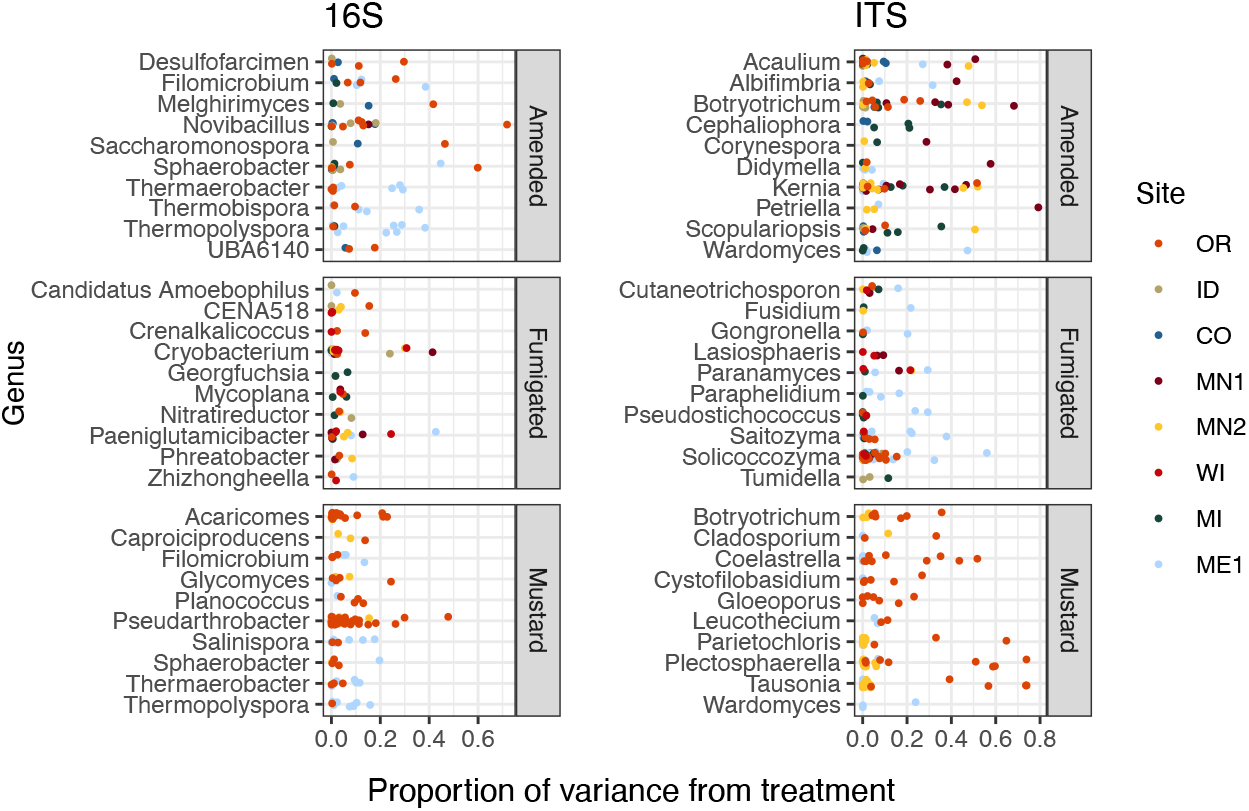
Proportions of variance in ASV relative abundance attributed to either amended, mustard, or fumigated soil treatments, for bacteria (left) and eukaryotes (right). ASVs are grouped by genus, showing the ten genera of each domain with the highest mean variance from each treatment, and which contained at least three ASVs across all field sites. Genera are sorted by alphabetical order.

Next, we used robust linear regression to associate ASV abundances with total tuber yields at harvest. *Yield-associated ASVs*, which accounted for 13.2% of bacterial and 11.9% of eukaryotic ASVs, were those whose relative abundances were significantly correlated with tuber yields (either positively or negatively, defined by p < 0.05 and R^2^ ≥ 0.2) at either microbiome sampling point during the 2022 season. Numbers of yield-associated ASVs varied idiosyncratically–without particularly noticeable trends–across rotation length, sampling time, and field site, but the distributions of R^2^ values from linear models remained broadly consistent, suggesting that overall, ASV associations with tuber yields were not discernibly different between planting or mid-season sampling points, two- or three-year rotation lengths, or bacteria and eukaryotes (Fig. S1).

Similarly to the patterns we observed in treatment-ASV associations, ASVs from the same genus displayed widely-varying correlations to total tuber yields (Fig. 3). Among the twenty most highly yield-associated bacterial and eukaryotic genera, most individual ASVs were not strongly positively or negatively associated with yields–rather, genus-level yield associations were strongly driven by comparatively few outlier ASVs with strong positive and/or negative correlations. Some genera included both yield-positive and yield-negative ASVs, sometimes within the same field site (e.g. *Actinoallomurus*, 16S, OR; *Rotundella*, ITS, ID). Consistent, genus-level directional associations to yields were not found across the continental US, but within individual field sites, genus-level directional associations with yield included the fungi *Chordomyces* and *Leptosphaeria* (ITS, positive, MN2), *Saitozyma* (ITS, weakly negative, ME1), and the bacterium *Nostoc* (16S, positive, CO; Fig. 3). Individual ASVs exhibited variable yield associations across field sites, rotations, and sampling times. Notably, a fungal *Microdochium* sp. included correlations to yield ranging from negative (p = 0.005, R^2^ = 0.28, MN1, two-year rotation, spring) to strongly positive (p = 4e-5, R^2^ = 0.50, CO, three-year rotation, spring) across field sites, rotation lengths, and sampling times (Fig. S2).

**Fig 3.**
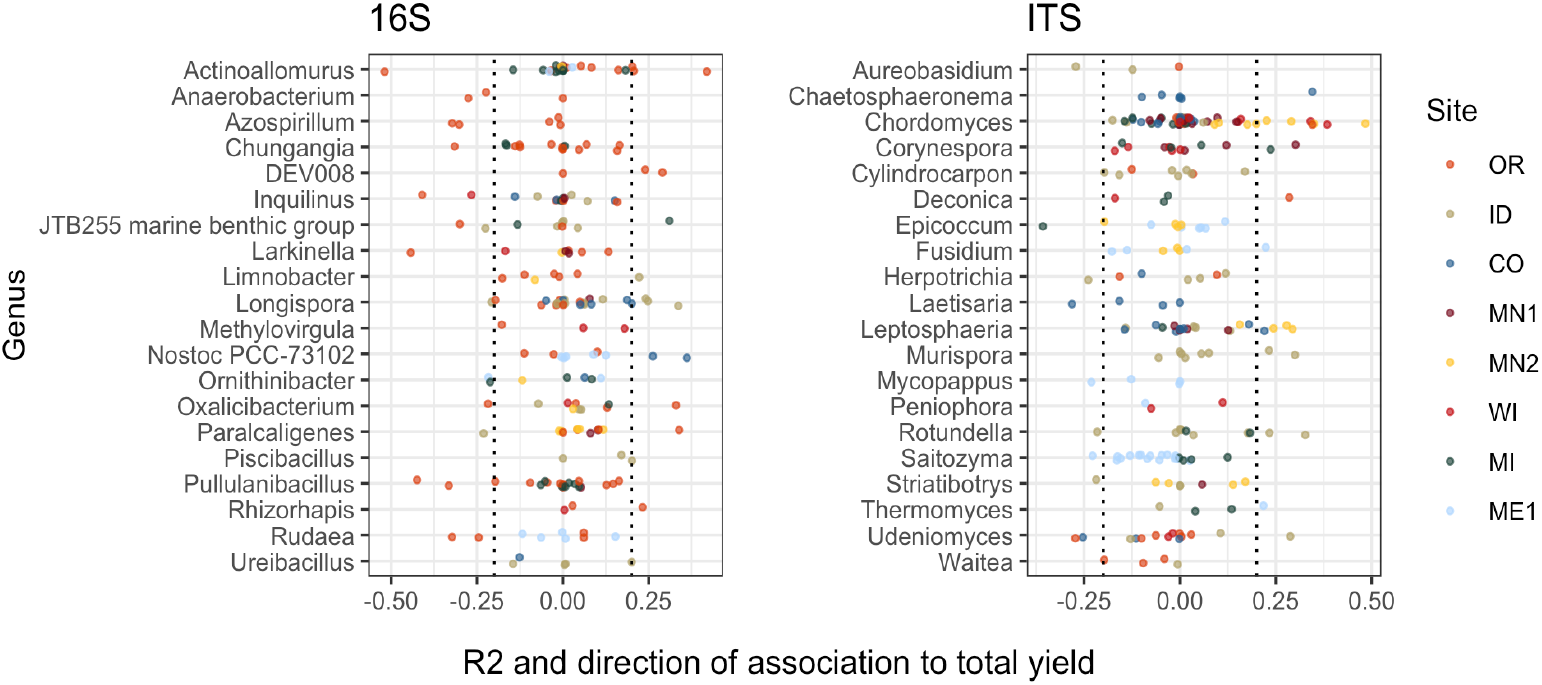
Bacterial and eukaryotic ASVs show variable associations with tuber yields. ASVs are colored by field site. X-axes show multiple R^2^ values of linear models with tuber yields predicted by CLR-transformed ASV abundance, for each combination of field site, rotation length, and sampling time. R^2^ values are multiplied by the sign of the association to yield (positive or negative). ASVs are shown for twenty genera from each amplicon with the highest mean R^2^ values (before accounting for sign of association) and at least three ASVs. Only ASVs present in at least 50% of plots within any field site, rotation length, and sampling time were modeled for relationships to yield. Dotted lines at ± 0.2 indicate correlation cutoffs deemed to be sufficiently yield-associated. Genera are sorted by alphabetical order.

To identify the ASVs most strongly linked to soil treatments and tuber yields, we narrowed our focus onto 45 target ASVs (34 bacterial, 11 eukaryotic). These represent the complete set of ASVs that both: 1) increased in relative abundance in a particular management scenario (in which microbiomes from both sampling times were combined); and 2) were significantly correlated, positively or negatively, with tuber yields at either sampling time of any management scenario (Fig. 4). Target ASVs were found in 16 of 33 management scenarios across six of eight field sites, and included several members of the bacterial phylum *Actinobacteriota* and fungal class *Sordariomycetes*. Soil treatment categories were associated with different numbers and yield associations of target ASVs: while organic amendments enriched 34 ASVs, 33 of which were positively associated with tuber yields, five of seven fumigation-enriched ASVs and none of seven mustard-enriched ASVs were yield-positive (Fig. 4). Target ASVs were largely site- and treatment-specific, with only three present in multiple field sites and none associated with more than one treatment category.

**Fig 4.**
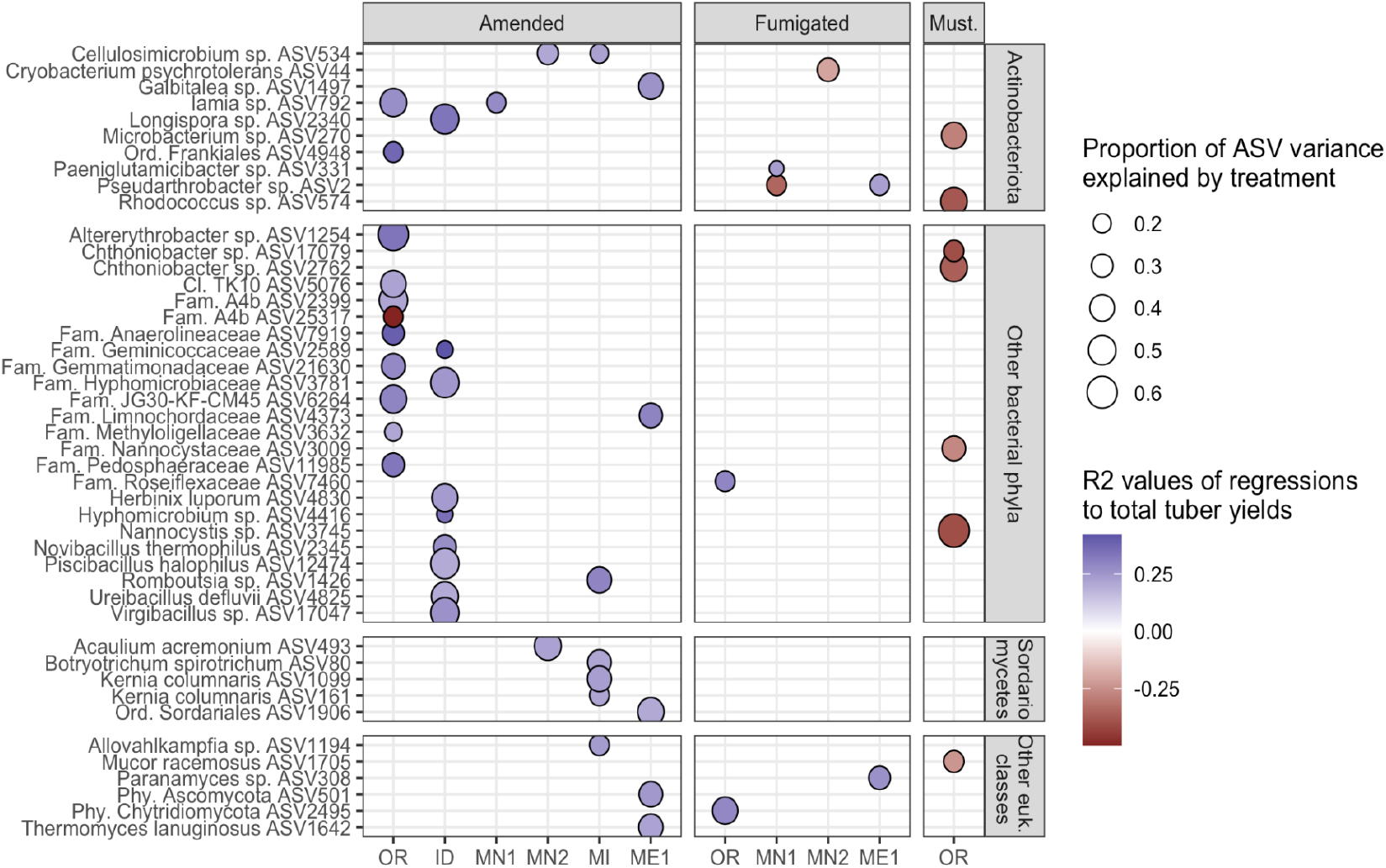
Target ASVs across field sites. All ASVs are enriched in amended, fumigated, or mustard-incorporated soil treatments relative to controls from the same field sites, with bubble sizes corresponding to the proportion of variance in ASV relative abundance explained by the treatment. Bubble colors represent R^2^ values of correlations to total yields; blue positive, red negative. The most specific taxonomic annotations to each ASV are given. Vertical panels are divided by domain (top two, bacteria; bottom two, eukaryotes). All eukaryotes here were classified as fungi except *Allovahlkampfia* (class *Heterolobosea*).

Once we identified target ASVs associated with 16 management scenarios across different treatments and field sites, we evaluated the predictive power of their cumulative abundances on total tuber yields at both sampling times during the 2022 season. At the same time, we leveraged a suite of soil biogeochemical data collected across all sites (pH, organic matter, ortho-phosphate, nitrate, ammonium, and bulk respiration) to evaluate whether yield increases could be attributed to changes in soil biogeochemistry associated with soil treatments, independent of shifts in microbiome composition. Among these 16 management scenarios, we identified a total of five microbiome-informed models (linear mixed models or structural equation models) in which abundances of target ASVs in bacterial or eukaryotic soil microbiomes could predict changes in yields where soil treatments or biogeochemical parameters could not. Target ASVs stimulated by organic amendments were associated with yield increases ranging from 10.4% to 23.2% in three models. In contrast, the only fumigation-based model (implicating one fungal ASV in OR) was associated with a modest 2.6% yield increase, and the only model involving mustard incorporation (on several bacterial ASVs, also in OR) showed a yield *decrease* of 26.4% (Fig. 5, Table 1, Supplemental Table S2). Interestingly, these latter two models applied to management scenarios that resulted in no significant changes to tuber yields; in other words, these treatments did not affect yields overall, but nevertheless influenced abundances of yield-associated taxa. These findings suggest the presence of other unidentified microbiome or soil chemical factors that may directly or indirectly counteract the predictive effects of the ASVs in the models. Interpreting these models as “false positives” (because the scenarios did not affect yields overall) suggests that this approach comprehensively identified the ASVs most strongly linked with treatments and yields. In eleven scenarios where target ASVs were identified, links between soil treatment, ASV abundance, and tuber yields were found to be below thresholds of statistical significance. Where no target ASVs were identified, no microbiome-informed models could be built (Fig. 5). Full details of these models are provided in Supplemental Table S2.

**Table 1.** All management scenarios where ASVs whose abundances were associated with significant changes in total yields relative to controls. Soil chemical parameters that changed relative to controls are listed, as well as model statistics and magnitudes of yield changes. Full model details are provided in Supplemental Table S2.

| Field site | Rotation length | Treatment | Treatment effect on total yield relative to control | Increases in soil chemistry | Decreases in soil chemistry | Model type | p-value | Multiple R<sup>2</sup> | Yield increase from target ASVs (Mg/ha) | Percent yield increase from target ASVs |
| --- | --- | --- | --- | --- | --- | --- | --- | --- | --- | --- |
| ME1 | 2 yrs | Amended | Increase | none | none | lm | 0.001 | 0.530 | 3.99 | 21.3 |
| OR | 2 yrs | Amended | Increase | %OM, pH, P, Nitrate-N, P, bulk respiration | NH<sub>4</sub>-N | lm | 0.005 | 0.728 | 14.50 | 23.2 |
| ID | 3 yrs | Amended | Increase | %OM, pH, P, bulk respiration | none | SEM | 0.275 | 0.614 | 6.57 | 10.4 |
| OR | 3 yrs | Fumigated | No effect | none | none | lm | 0.013 | 0.609 | 1.78 | 2.6 |
| OR | 3 yrs | Mustard | No effect | Nitrate-N | pH | SEM | 0.099 | 0.668 | -17.63 | -26.4 |

**Fig 5.**
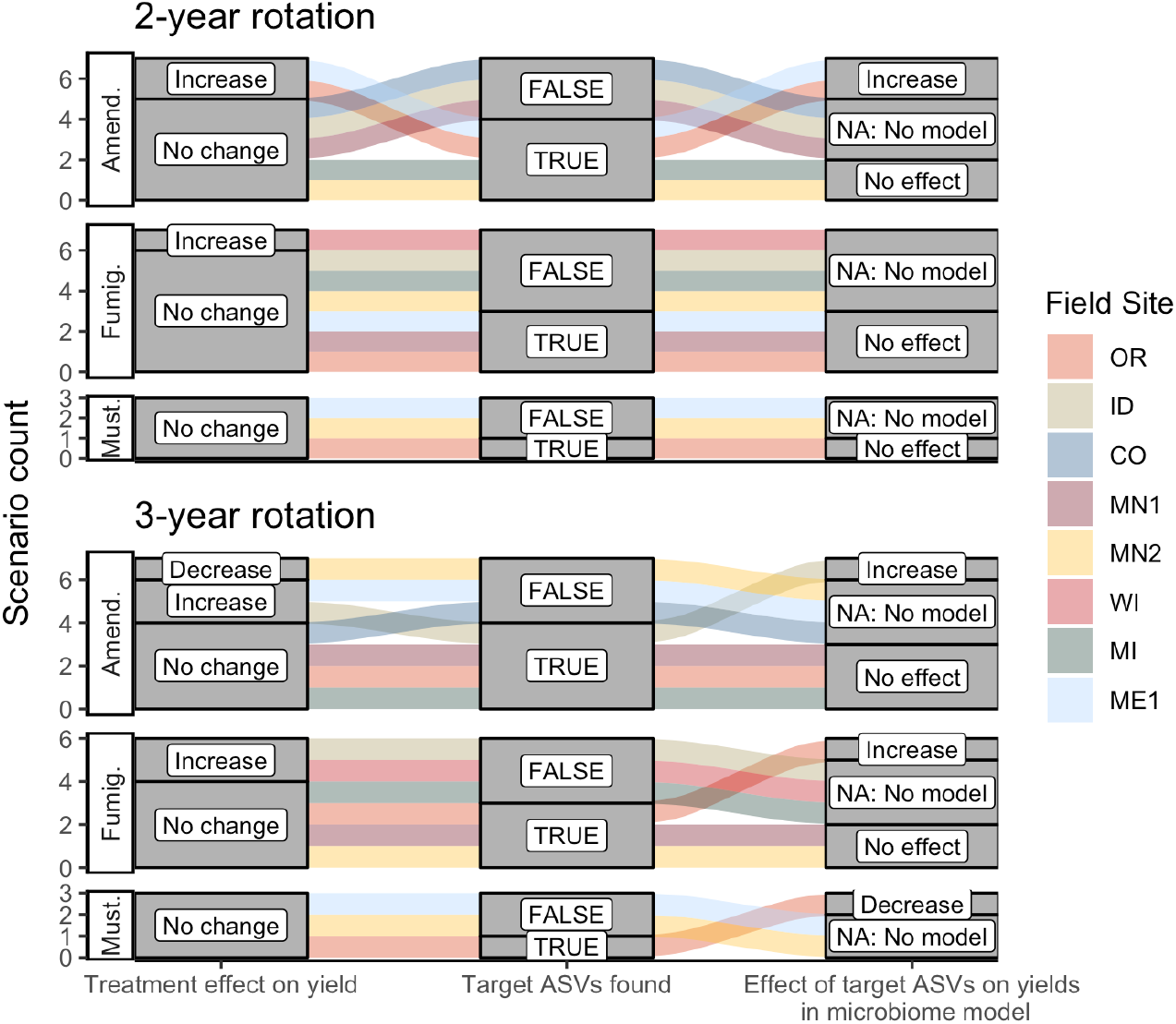
Alluvial plots depicting the detection and contribution of target ASVs towards tuber yields in each management scenario. The left columns indicate whether any significant change in total tuber yield was observed within the dominant cultivar grown at each site, relative to corresponding untreated plots. The middle columns indicate whether target ASVs were detected in bacterial or eukaryotic microbiomes at either timepoint during the 2022 sampling (ASVs shown in Fig. 4). The right column shows whether increased abundances of target ASVs in treatments (relative to non-treatment controls) could explain increases or decreases in total yields within microbiome-informed models. Note that no models could be constructed when no target ASVs were found. Treatments: Amend., Amended; Fumig., Fumigated; Must., Mustard.

None of the organic amendment-stimulated ASVs from any microbiome model belonged to the same genus, highlighting the narrow taxonomic specificity of these treatment-microbiome-yield relationships. However, microbiome-informed models of amendments in OR and ID field sites were noteworthy in that they linked the compost amendment-associated stimulation of several bacterial target ASVs to increases in tuber yields (a similar model for ME1 implicated two fungal ASVs). While no target ASVs were shared across models for OR and ID sites (Fig. 4, Supplemental Table S2), the same ASVs were present–though not yield-associated–in soil microbiomes from fields in both states. In fact, compared to plots sampled in 2019 or 2020, all nine target ASVs from the ID 3-yr rotation amendment model were also stimulated by amendments in the OR field site to nearly equal proportions within bacterial microbiomes (Fig. 6A). It is interesting to note that while these microbiome shifts accounted for a 10.4% increase in yields in ID, in total they represented only a modest proportion (∼0.4%) of soil bacterial microbiomes by abundance, even after enrichment.

**Fig 6.**
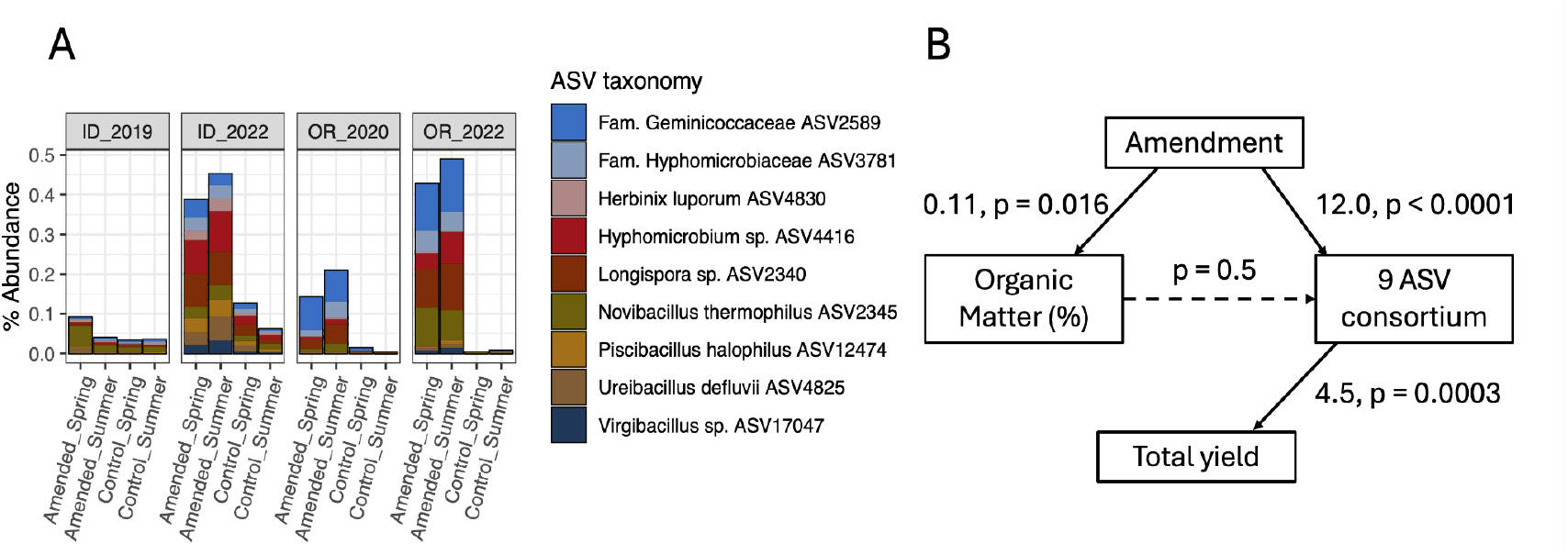
Compost amendments stimulate a consortium of nine bacterial ASVs in OR and ID field sites that are associated with increased yields only in ID. **a** Mean percent abundances of nine amendment-stimulated ASVs within bacterial communities across replicate (n = 4) plots in ID and OR field sites in potato growing years (ID 3-yr rotation to 2019 and 2022; OR 2-yr rotation to 2020 and 2022, separated by soil treatment and timing of sampling). **b** Structural equation model (SEM) showing relationships between compost amendment (coded as a binary dummy variable), soil percent organic matter, CLR-transformed sum relative abundances of the nine bacterial ASVs, and total tuber yields in Idaho 2022 samples (summer 3-year rotations, Norkotah cultivars only, n = 14). Linear model estimates and p-values are shown for each path, with significant paths represented by solid arrows.

Two of the three organic amendment scenarios that resulted in yield increases and significant microbiome-informed models also changed soil chemistry: percent organic matter (OM), pH, and ortho-phosphate increased in both ID and OR, while OR soils additionally increased in nitrate and bulk respiration, and decreased in ammonium (Table 1) in response to compost amendment. Additionally, the mustard amendment in OR associated with yield-negative ASVs decreased soil pH and increased nitrate. However, as illustrated in the model of amendment in ID (Fig. 6B), none of the soil biogeochemical changes in any of these models directly influenced ASV abundances. These key taxa may thus operate independently from–or tangentially related to–management-caused changes in soil chemistry.

## Discussion

Amidst a growing recognition of the many ways in which soil microbiomes support plant health and productivity, we leveraged a continental-scale platform consisting of soil chemical, tuber yield, and microbiome data from eight field sites in potato growing regions of the US to investigate how soil management practices used in potato production affect soil microbiomes and tuber yields. We specifically asked 1) Whether soil treatments affect soil microbiomes in consistent ways across diverse growing regions; and 2) Are distinct microbial taxa or consortia linked with soil treatments and tuber yields across regional or continental scales? We address these questions after first providing context on how soil treatments affected yields across the continental-scale platform.

### Experimental design, treatment variability, treatment effects on yields

All field sites implemented a four-year rotation with potatoes grown at both two- and three-year intervals, in a factorial design with six discrete treatments. Selection of cultivars and cover crops as well as composition of treatments within each field site were chosen to reflect regionally-specific growing practices. Treatment conditions, such as types and application rates of fumigants or organic amendments, varied widely; compost amendment rates of 25 to 37 metric tons/hectare, for example, were designed to characterize the upper bounds of soil chemical and microbial changes. This treatment heterogeneity, particularly in the type and amounts of organic amendments applied across field sites, may partially account for variable influences on yields (Fig. 1B-C, Fig. 5). This is consistent with prior observations that different types of organic amendments exert variable, but generally positive, effects on crop yields (32). Differences in soil texture, soil chemistry, and irrigation status, such as what we previously observed in these field sites (16), can also influence the extent to which yield increases are observed with organic matter application (32). Regionally variable effects of metam sodium fumigation on tuber yields across growing sites (Fig. 1B) have also been noted on grower fields in Wisconsin, highlighting complex relationships with soil texture, chemistry, and microbiome diversity (33).

The four-year timescale of this study is another important factor when considering that significant changes in yields, soil health, and microbiomes may require consistent, repeated treatments across longer timescales. For example, decadal-scale implementation and monitoring of potato soil treatments has revealed changes in soil chemistry and disease suppression that can take several years to develop (8). The efficacy of organic amendments on tuber yields did not seem to relate to prior history of potato cultivation, as fields that were both previously potato-naive (OR and CO) and fields that had a history of potato prior to 2019 (all others) each showed mixed results. Nevertheless, this dataset, which consists of 96 specific soil treatments from eight field sites across the continental US (Supplemental Table S1), provides the most comprehensive and agriculturally-relevant picture of how potato yields and soil microbiomes respond to diverse management practices.

### Treatment effects on microbiomes

We did not observe regionally consistent patterns in how potato soil microbiomes were affected by fumigation, organic amendment, and mustard incorporation treatments (Fig. 1C). The variable influences of these treatments on soil microbiomes across field sites likely reflect regional- and continental-scale variation in soil texture, chemistry, and initial microbiomes themselves, particularly at genus and finer-scale taxonomic levels (16). Specific implementations of each treatment, such as the type, amount, frequency, and timing of application likely contribute to their varying influences on soil microbiome composition across field sites. The overall low impact of metam sodium fumigation on soil microbiomes–particularly bacterial ones–may reflect the fact that all fumigation occurred in fall 2021 prior to planting in spring 2022. Metam sodium transiently alters bacterial community composition in the weeks following application, after which the effects of fumigation tend to subside (34,35). In OR, the strong influence of mustard incorporation on bacterial and eukaryotic microbiomes (Figs 1C, 2) may reflect the fact that mustard was planted and tilled into plots every year of the four-year sequence, in contrast to one- or two-year applications at other field sites (Supplemental Table S1). Soil management, and organic amendments in particular, can influence community assembly patterns of soil microbiota (36). Variable effects of organic matter addition on microbiomes may be linked to the complex ways in which they interact with bacterial and fungal phyla relating to soil texture, climate, amendment type, and nutrient stoichiometry (37,38). It is possible that the comparatively high variance attributed to rotation length, particularly for eukaryotic microbiomes, is linked to differences in disease pressures and cropping history between rotation cycles, as noted previously (39,40). Nevertheless, further work is needed to untangle the complex biotic, abiotic, and crop management history interactions that mediate short-term (1-4 year) responses of soil microbiomes to agricultural treatments.

### Treatment effects on ASVs, ASVs and yield, heterogeneity therein

We did not identify consistent correlations between microbial taxa (either bacteria or eukaryotes, at the ASV or genus levels) and soil treatments or tuber yields across regional scales. Instead, our finding that distinct ASVs belonging to the same bacterial or fungal genera were impacted by soil treatments to widely varying degrees (Fig. 2) suggests that closely-related ASVs may respond quite differently to certain soil treatments, even within the same field site. Differential responses among specific members of soil microbial populations may reflect both ecological and evolutionary processes, as recently demonstrated in a grassland climate gradient (41). It is plausible that four years of applied soil treatments–particularly ones that increase soil organic matter or reduce pathogen populations–impart selective pressures among members of microbial populations to drive within-field-site differentiation across treatments, similarly to the differentiation across field sites we previously described (16).

Associations between ASV abundances and tuber yields (Fig. 3) can be interpreted in many ways. With this dataset, it is not possible to discern whether these associations reflect direct plant-microbe interactions that promote or inhibit plant growth (such as facilitating plant nutrient uptake, or pathogenicity), indirect interactions (pathogen suppression, or inhibition of plant growth-promoting microbes), or unrelated responses to the same stimulus (e.g., organic amendment independently increasing both tuber yields and abundances of certain ASVs). In root-associated bacteria, horizontal gene transfer has distributed genes contributing plant-beneficial functions across wide taxonomic ranges (42,43), and recent evidence suggests that key functional traits can play a larger role than taxonomy in predicting plant growth responses (44). Thus, within field sites, we attribute genus-level variability in yield associations to species- or strain-level differences in functional capacity; across field sites, additional variability may be driven by differences in edaphic or climatic factors, management history, and microbiome composition (16). An implication of this variability is that the identification of soil bacterial or fungal populations or functions as potential indicators of soil health and tuber yields may be most likely relevant at regional or local (rather than continental) scales, likely in conjunction with other key contextual information such as soil chemistry, rotation length, or type of amendment.

Within this amplicon dataset, modeling associations at the ASV level was critical for uncovering fine-scale taxonomic variation in treatment- and yield-associations. While this allows for the possibility that a clonal strain with multiple amplicon copy numbers may be represented by multiple distinct ASVs, investigating patterns at this level (as opposed to higher taxonomic categories, e.g. OTU or genus) can enhance predictive power (14,45), and additionally identify strain-specific responses to treatments (46).

### Microbiome models

Although we could not identify clear patterns between microbial taxa and soil treatments or between taxa and tuber yields, we were able to identify a set of 45 target ASVs associated with both treatments and yields across field sites (Fig. 4). This permitted us to explicitly model whether soil treatments increased relative abundances of any of these taxa in ways that might account for significant changes in tuber yields, relative to non-treatment control plots (Fig. 5, Supplemental Table S2). Modeling specific treatment scenarios from this continental-scale dataset required subsetting samples by field site, treatment, rotation length, and sampling time, due to the limited occupancy of ASVs across field sites (16) and the variability in ASV-yield associations across time and rotation length (Figs. S1 & S2). While this reduced sample sizes to 12 to 25 measurements (of microbiome taxa abundances, tuber yields, and soil chemical data) per model, the high number of field sites and soil treatments in this dataset allowed us to obtain a robust picture of how commonly-used soil treatments across US potato growing regions affect soil microbiomes and their relationships to tuber yields and soil chemistry.

Compared to fumigation and mustard application, organic amendments were more frequently, and more strongly, linked with microbiome-associated increases in tuber yields. Inconsistent relationships between fumigation, yields, and soil microbiome characteristics have also been attributed to differences in microbiome diversity or microbial carbon cycling potential across growing sites with distinct soil chemical profiles (33). At global scales, soil organic amendments shape soil physical properties and increase microbial biomass, soil microbial C/N/P acquisition, and cereal crop yields (32,47). Organic amendments in potato soils have been linked to enrichment of soil microbial taxa commonly implicated in pathogen suppression (15), notably within the ME1 field site (Ashley et al, in review).

Two interrelated remaining questions are 1) why significant microbiome-informed models could be built for only three of fourteen organic amendment scenarios, and 2) for field sites where we identified such models, why we observed inconsistencies between two- and three-year rotation lengths. A possible, partial explanation for the first question may involve the particularly high input rates of compost applied to these treatments (ID and OR field sites used 10 and 15 T/ac respectively). As for the second question, we observed that higher microbial biomass and fungi:bacteria ratios in ID and OR field sites coincided with rotation lengths where compost amendments stimulated microbiomes to increase yields (Fig. S3), which may support the idea of a critical soil microbial biomass threshold required for treatments to be effective. Interestingly, we did not find that soil percent organic matter was linked to abundances of target ASVs in microbiome-informed models for either ID or OR field sites (Fig. 6B, Supplemental Table S2), which potentially discounts the role of organic matter as a direct nutrient source for enriching target ASVs and stimulating tuber yields. Instead, organic matter application may act to shift nutrient utilization and inhibitory behaviors of key soil taxa (48). Finally, the discrepancy in yield associations in the consortia of organic amendment-enriched bacterial ASVs from ID and OR (Fig. 6A) suggest differences in functional potential across field sites from the same growing region, justifying further efforts to characterize functional variability of plant-health-associated microbiota (particularly as potential inoculants) across agricultural contexts (49).

### Future directions

While the staggering complexity of soil microbiomes presents challenges for predicting microbial functions from amplicon sequence data (50), recent efforts have yielded a molecular index predictive of several commonly-used soil health indicators across diverse soil types (51). Characterizing specific mechanisms of action by which soil amendments increase abundances of certain taxa, and by which these taxa lead to higher tuber yields, is beyond the scope of this study. But for specific scenarios where these effects were detected, temporally-resolved sampling of soil chemistry, microbial genes and transcripts, and plant and microbial metabolites could be used to establish such relationships (2). Linkages between treatment-stimulated soil taxa and endophytic taxa may be of particular importance in potatoes, as machine learning models have recently used endophyte microbiomes to predict tuber yields in subsequent growing seasons (14). In addition, the largely microbiome-independent effects of fumigation on yields call for more thorough mechanistic investigations. While this study provides a crucial baseline for linking management practices to microbiome taxa and tuber yields across the continental US, efforts to integrate large-scale soil health and microbiome data using machine learning approaches are needed to reveal regional- or field-specific targets for managing soil and crop health (45).

## Acknowledgements

We thank Julia Ahlborn, Zoe Hansen, Aspen Hughes, Brett Lane, Matthew McNearney, and Seonghyun Seo for performing lab and fieldwork. Marian Bolton developed soil sampling protocols, assisted with fieldwork, and provided logistical support. Corbin Dirkx at the University of Minnesota Genomics Center oversaw and performed amplicon library preparation and sequencing. This work is an outcome of the Potato Soil Health Project (https://potatosoilhealth.cfans.umn.edu/) managed at the University of Minnesota.

