## supplemental_figures for "Compost amendments to potato soils enrich yield-associated members of soil microbiomes across the continental US"

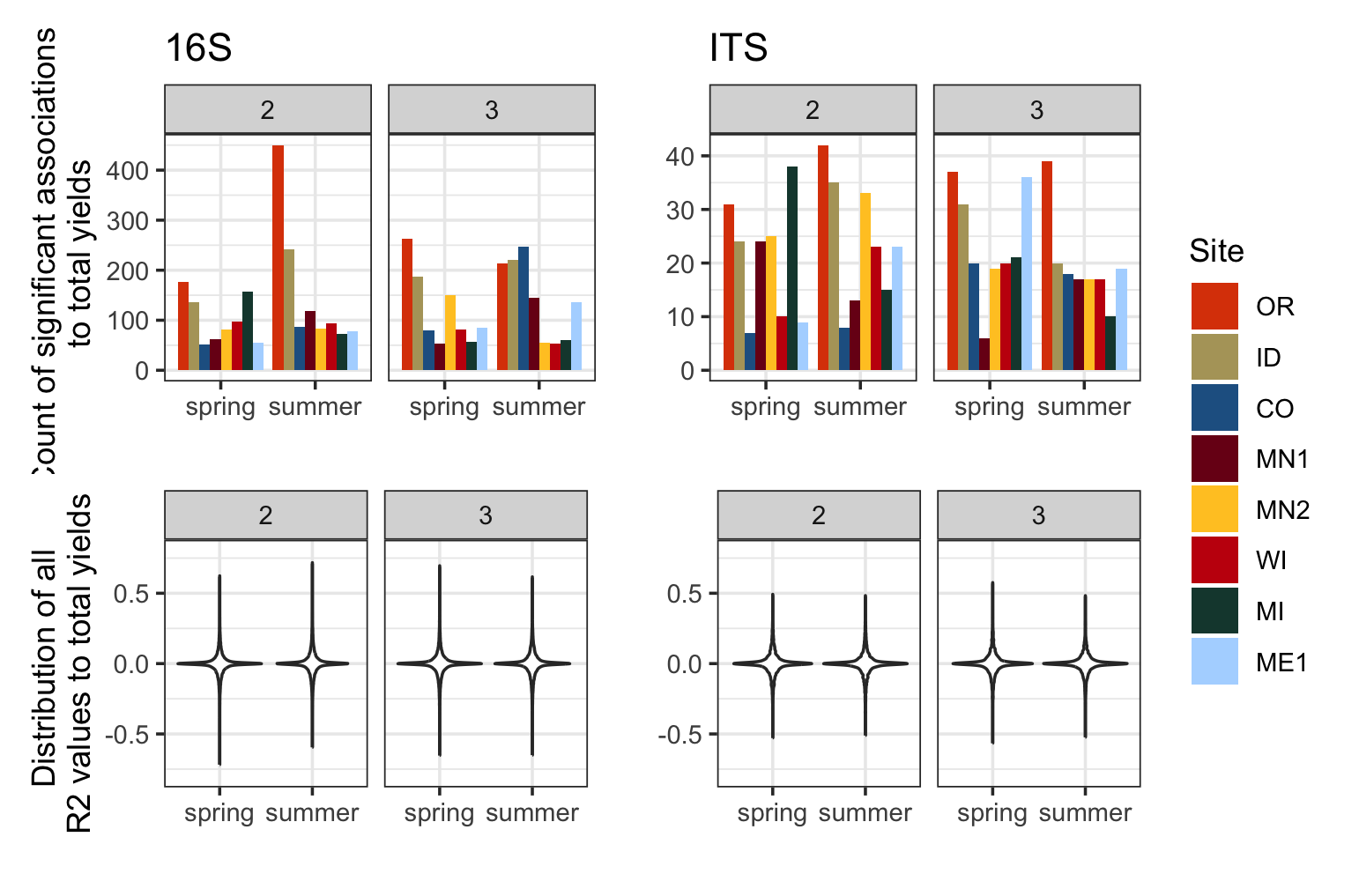

**Figure S1.** Numbers of significantly yield-associated bacterial and eukaryotic ASVs across field sites, rotation lengths, and sampling timepoints (top) and distributions of R^2^ values from each ASV-yield regression (bottom). R2 values are multiplied by the sign of associations to tuber yields. Spring and Summer time points correspond to “at planting” and “60 days after planting”, respectively.

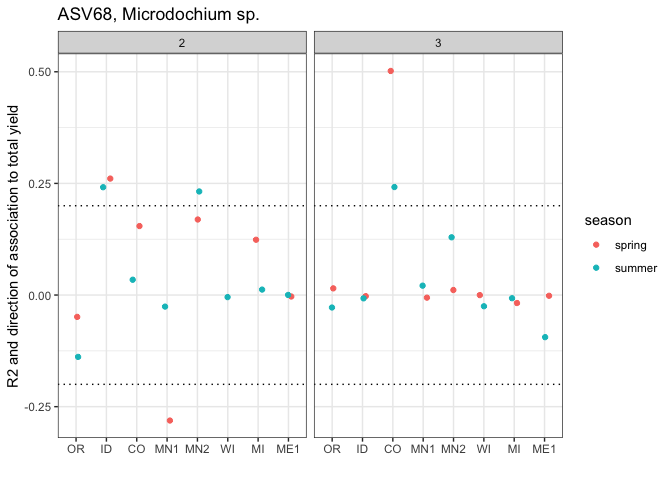

**Figure S2.** R^2^ values of positive and negative ASV-yield regressions across field sites, rotation lengths, and sampling times for fungal ASV68, a *Microdochium* sp. Dotted lines at $\pm$ 0.2 indicate correlation cutoffs deemed to be sufficiently yield-associated. Spring and Summer time points correspond to “at planting” and “60 days after planting”, respectively.

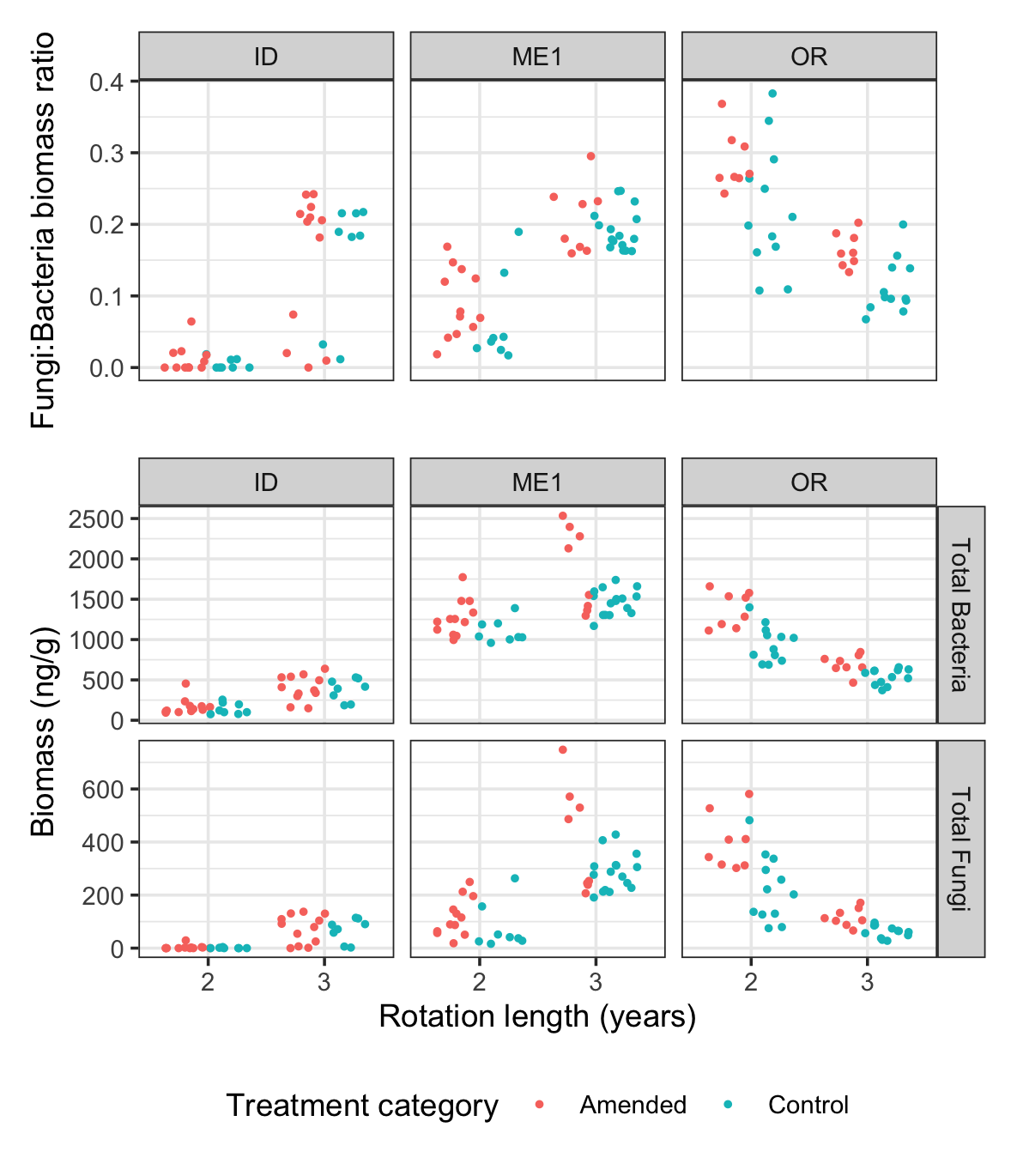

**Figure S3.** Summer 2022 PLFA measurements of soil bacterial and fungal biomass, and bacterial/fungal biomass ratios in ID, ME1, and OR, by rotation length and treatment.

### Supplemental Tables

Table S1. Management details and four years of cropping history for all soil treatments applied at each field site.

Table S2. All management scenarios listed by effects of treatment on yield, and whether target ASVs were detected. Where relevant, details of microbiome models built from scenarios are provided, along with ASVs and soil chemical data used in models.
